# TFAM organizes DNA into compact higher order structures

**DOI:** 10.64898/2026.04.24.720727

**Authors:** Sashi R. Weerawarana, Wei Tian, Karolin Luger

**Affiliations:** Department of biochemistry, University of Colorado, Boulder, CO, USA; Howard Hughes Medical Institute, Chevy Chase, MD, USA

**Keywords:** TFAM, mitochondria, cryo-EM, mass photometry

## Abstract

TFAM (Transcription Factor A, Mitochondrial) is an essential human protein that plays two key roles in mitochondrial DNA (mtDNA) homeostasis. TFAM acts as a transcription factor that specifically binds to promoter regions, but it is also solely responsible for organizing mtDNA into nucleoids by nonspecifically covering the entire genome. Many studies have addressed TFAM in transcription regulation, but its role as a genome organizing entity is not well characterized. The current understanding of how TFAM compacts DNA into nucleoids is based on crystal structures of a TFAM monomer bound to short fragments of DNA (22-28 bp). However, this does not adequately reflect the biological role of TFAM in organizing the nucleoid where multiple TFAM molecules oligomerize on the 16.5 kb genome to form the nucleoid. Here, we present a biochemical and structural analysis of TFAM oligomerization on longer DNA. Our results show that TFAM compacts longer segments of DNA into higher order complexes that are homogenous yet exhibit continuous conformational dynamics.

**Significance statement:** Mutations or damage to mitochondrial DNA (mtDNA) severely impairs cellular respiration and is implicated in many human diseases and aging. ‘Transcription Factor A, Mitochondrial’ (TFAM) is an essential protein that is solely responsible for packaging mtDNA into nucleoids thereby shielding it from DNA damage. Despite its importance, the mechanism by which this is accomplished is poorly understood. Here, we use biochemistry to show that TFAM oligomerizes on DNA to form compact, homogenous higher order structures in solution. We also examined these complexes at a low resolution using cryo-EM, suggesting an organizational unit of mtDNA. This work reveals there may be a regular organization to mitochondrial nucleoids, providing the basis for further understanding mtDNA compaction by TFAM.

## Introduction

Human mitochondria contain a 16.5 kb circular genome that encodes 13 essential protein subunits of the electron transport chain, along with several essential RNAs^1^. Mutations or damage to mitochondrial DNA (mtDNA) is implicated in many diseases and contributes to aging^2–4^. Within mitochondria, mtDNA is packaged by the protein TFAM (Transcription Factor A, Mitochondrial) into complexes termed nucleoids that are ∼100 nm in length. A single mitochondrion contains multiple nucleoids, each consisting of a single copy of mtDNA coated by ∼1000 TFAM molecules^5,6^.

TFAM is an essential, nuclear-encoded HMG box (high mobility group box) protein consisting of two HMG domains separated by a linker helix. TFAM plays two important roles in mtDNA homeostasis, by functioning both as a transcription factor and a genome organizer^1^. First discovered as a transcription factor^7,8^, TFAM specifically binds to the promoter regions of mtDNA to recruit the mitochondrial polymerase, POLRMT, and a second transcription factor, TFB2M, to form the preinitiation complex^9,10^. High resolution structures of the mitochondrial preinitiation complex demonstrate that TFAM induces a sharp U-turn distortion in the DNA that is required for the recruitment of the transcription machinery^9,10^. Each HMG box binds to the minor groove of the DNA through numerous interactions between basic side chains and the DNA backbone, and a conserved leucine side chain that intercalates in the minor groove to introduce a 180° bend in the DNA^11–15^.

It was later discovered that TFAM is also the sole protein responsible for compacting mtDNA into nucleoids^6,16^. mtDNA accessibility is regulated by the ratio of TFAM to mtDNA^17,18^. The majority of nucleoids in a given mitochondrion exist in a genetically silent state where they are fully coated and compacted by TFAM molecules^17,18^. It is hypothesized that these inaccessible nucleoids are shielded from DNA damaging agents, serving as a genetic reservoir for undamaged mtDNA. Despite the importance of mtDNA packaging by TFAM, the molecular details of how TFAM oligomerizes on and compacts the mitochondrial genome remain unknown.

Because all available crystal structures of TFAM were done in complex with short 22-28 bp fragments of DNA^11–15^, it is currently unknown how TFAM oligomerization contributes to DNA compaction. TFAM titration onto plasmid DNA visualized with low resolution methods (rotary shadowing EM and Atomic Force Microscopy) show evidence of higher order structures formed by TFAM multimers^5,19^. However, the details of these higher order assemblies cannot be determined at these resolutions.

Here we conduct a biochemical and structural analysis of TFAM-DNA complexes utilizing fragments of DNA that are long enough for multiple TFAMs to associate. We sought to determine if there is any regular manner in which TFAM oligomerizes on DNA, or if mtDNA is compacted by the random association of TFAM with no ordered higher-level organization.

## Results

### Multiple TFAM molecules compact DNA into complexes that are uniform in size and shape

DNA fragments of varying lengths (80 bp, 160 bp, 240 bp and 400 bp) of a randomized sequence were generated (see methods) to assess TFAM binding and oligomerization. We chose to use a randomized sequence to characterize TFAM interaction with “average” DNA and prevent any sequence bias. All DNA lengths used are long enough to accommodate multiple TFAM molecules, assuming a footprint inferred from published crystal structures (∼22-28 bp per TFAM). The ability of TFAM to bind these DNAs was validated using Fluorescence Polarization (FP). Consistent with published data, TFAM bound all DNA fragments tested here cooperatively with EC_50_ values in the nanomolar range^11,19–22^ (**Figure S1**). The 10-fold difference in affinity between the 80mer and 400mer is likely due to the larger number of binding sites on the 400mer.

DNA binding and stoichiometry was further assessed using Electrophoretic Mobility Shift Assays (EMSAs) and mass photometry. Although TFAM binding is highly cooperative (Hill coefficients > 1.5 in binding experiments, Fig. S1), EMSA analysis showed a distinct laddering pattern consistent with multiple discrete binding events at lower ratios, especially for the shorter DNA fragments (**Figure 1A**). To monitor TFAM oligomerization in solution, the same samples were analyzed by mass photometry. This is a powerful in-solution technique that provides a molecular weight distribution of species in a given complex. We obtained molecular weights for each species for each of the ratios, and from that calculated the number of TFAM molecules bound to the DNA. Using this method, we determined the saturating TFAM:DNA ratio for each DNA length based on the maximum molecular weight observed on the mass photometer upon further TFAM titration (**Figure 1B, Figure S2**). This allowed us to calculate a footprint for TFAM (i.e. number of base pairs occupied by each TFAM molecule). For the four DNA lengths tested, the saturating ratio yields a homogenous complex with a calculated footprint of ∼15 bp/TFAM (**Figure 1B**). This is in contrast with crystal structures of TFAM on shorter DNA in a 1:1 complex, which consistently suggest a footprint of ∼22-28 bp/TFAM^11–15^. The near-identical footprint observed across different lengths of DNA implies that TFAM likely adopts the same binding mode in each complex. Importantly, our mass photometry data also shows that TFAM forms saturated complexes that appear to be homogenous in size on DNAs as long as 400 bp (**Figure 1B, Figure S2**).

**Figure 1.**
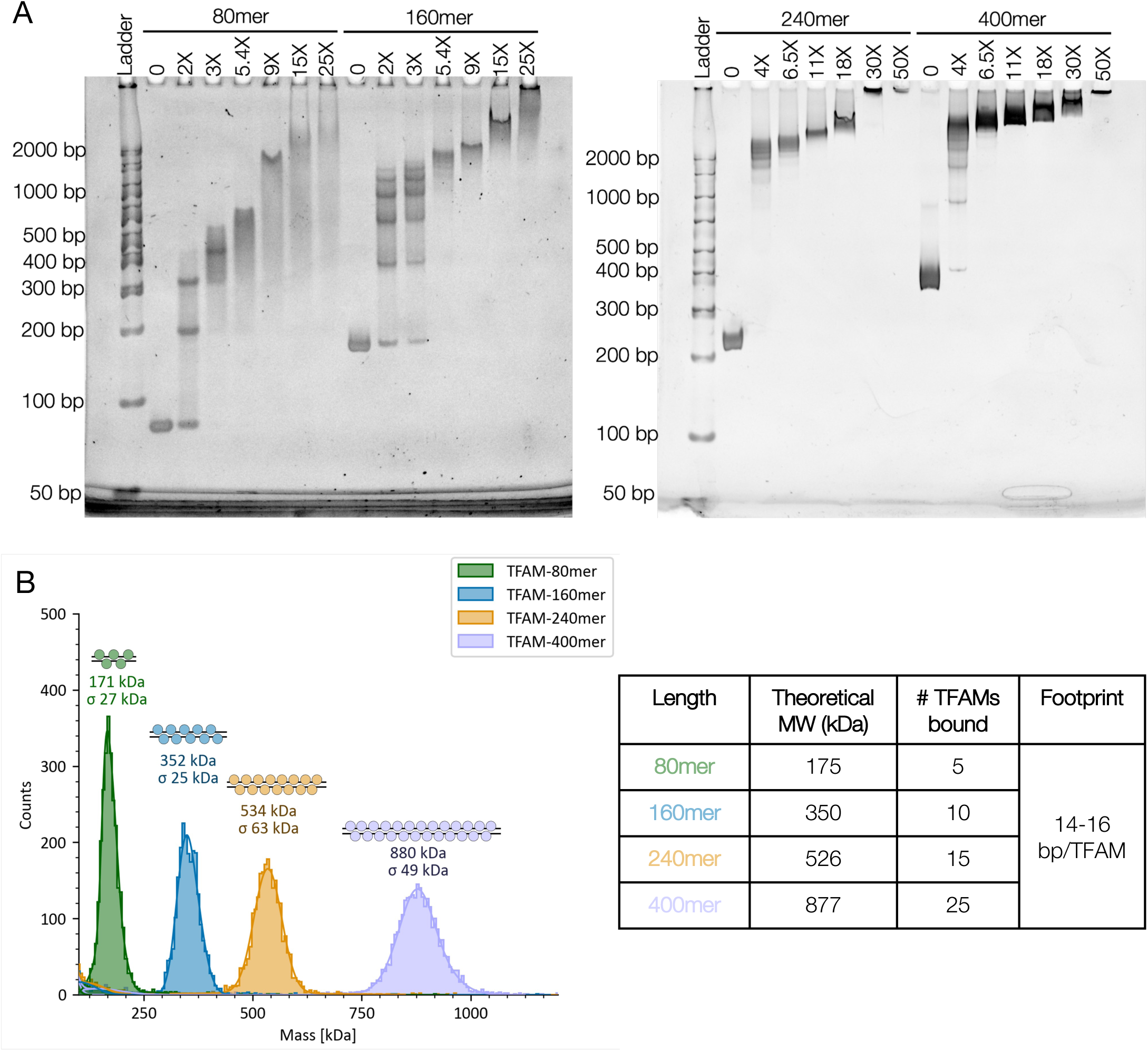
Multiple TFAM molecules compact DNA into homogenous complexes. **A.** EMSA analysis of TFAM titration into a fixed concentration of the indicated DNA lengths at a DNA concentration of 1 µM. Molar equivalence above each lane indicated-fold excess of TFAM over DNA molecule. Samples were run on a 6% polyacrylamide gel (Invitrogen) and stained with ethidium bromide. **B.** Mass photometry of saturated complexes with the indicated DNA lengths. The number of TFAM bound, and TFAM footprint on the DNA calculated from the molecular weights are shown in the inset. Data for each titration point are shown in Figure S2. For both EMSA and mass photometry data, n=3 for 80mer and 160mer; n=2 for 240mer and 400mer. Representative data is shown.

Analysis of the intermediate binding events under sub-saturating conditions by mass photometry showed that individual TFAM binding events could be detected using this method, particularly in the two shorter lengths (**Figure S2A-B**). This result contradicts the prevailing notion that TFAM binds as a dimer^12,19,21,23^, with the caveat that the low nanomolar concentrations used here may not reflect the actual binding mechanism. The low concentrations also explains why individual binding events were detected despite the steep Hill slope observed in FP experiments (**Figure S1**), where concentrations were two-fold higher.

While mass photometry can robustly determine the number of assemblies in a sample and their respective molecular weights, it does not provide any information on the shape of these complexes. To assess if TFAM-DNA complexes are homogenous in shape, we performed Sedimentation Velocity Analytical Ultracentrifugation (SV-AUC) with TFAM-160mer complexes at varying levels of saturation (**Figure 2**). SV-AUC uses first principles to directly assess the diffusion-corrected sedimentation (s) and frictional coefficient (f/f0) of a sample. The s-value provides information on size, while f/f0 describes the viscous drag of a complex, which conveys information on shape or compactness. The distribution of these values provides information on sample heterogeneity. Molecular weights can also be calculated from AUC; however these are not derived from first principles and should be validated using orthogonal methods such as mass photometry. We tested four TFAM-160mer complexes of varying saturation: 5X (undersaturated), 10X (saturated), 15X and 20X (oversaturated). Increasing the concentration of TFAM at constant DNA concentration increased sedimentation coefficients and decreased the frictional coefficient, consistent with further compaction of the DNA with increasing TFAM saturation (**Figure 2**, **Table 1**).

**Figure 2.**
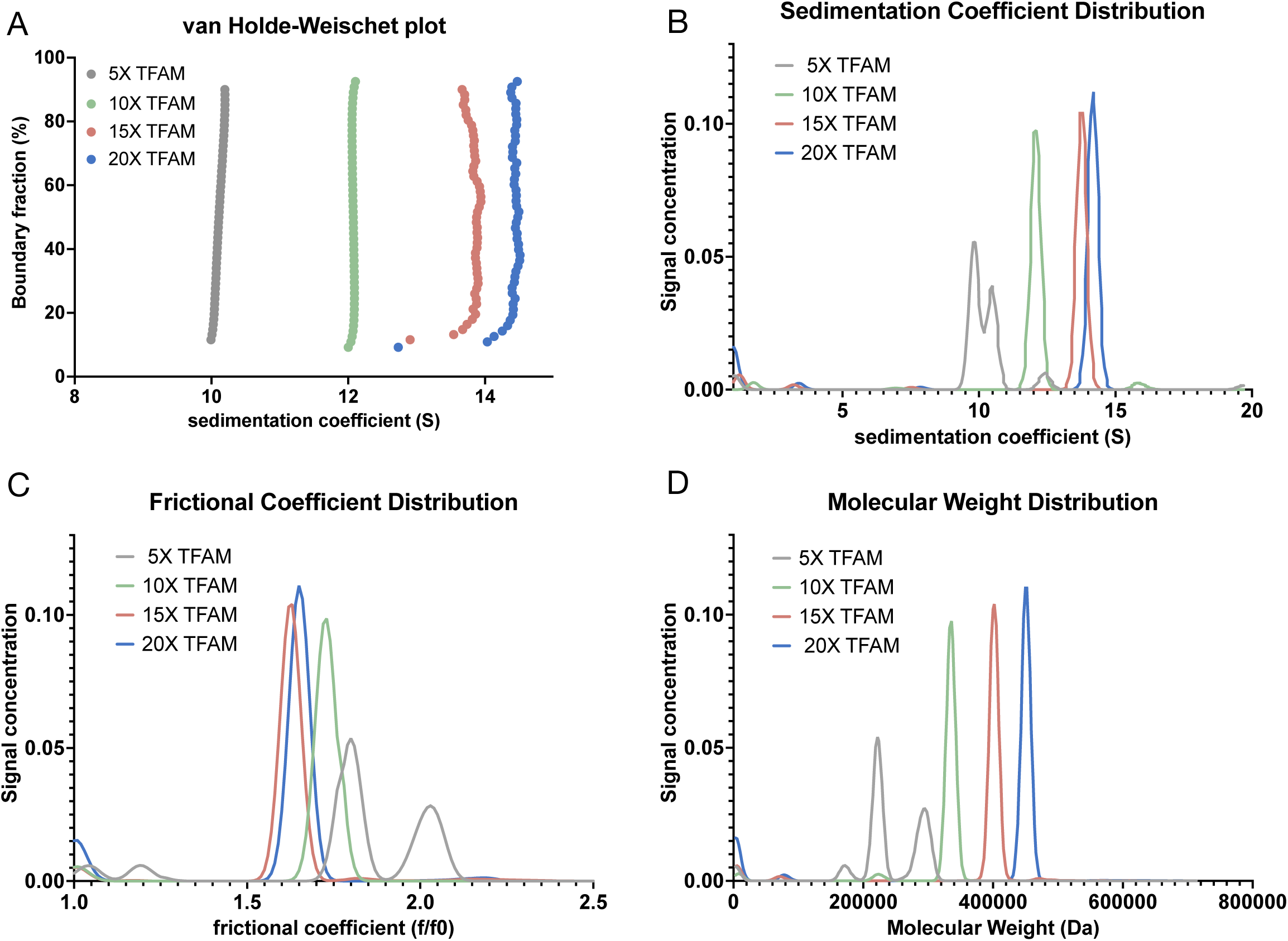
SV-AUC analysis shows that TFAM complexes adopt a compact shape. Four TFAM-160 bp complexes of varying saturation were tested: 5X TFAM (undersaturated), 10X TFAM (saturated), 15X and 20X (oversaturated). **A.** Van Holde-Weischet plot showing sedimentation (s) coefficients as a function of boundary fraction. **B-D.** Distribution plots showing s-values (B), frictional ratios (C) and molecular weights (D) as a function of signal concentration. All values were calculated using Ultrascan III25. Exact values for each complex and 95% confidence intervals can be found in table 1. n=1.

**Table 1.** AUC and mass photometry values of TFAM-160mer complexes at varying degrees of saturation. Table shows s-values, frictional ratios, molecular weights derived from AUC and molecular weights derived from mass photometry (MP). 95% confidence intervals are shown for s-value, f/f0 and AUC molecular weights. Standard deviation from a Gaussian fit are shown for MP molecular weights. AUC data are n=1 and mass photometry values are representative from n=3.

| TFAM equivalence | Sedimentation coefficient (s(20,w)) | Frictional ratio (f/f0) | AUC molecular weight (kDa) | MP molecular weight (kDa) | Theoretical MW (kDa) |
| --- | --- | --- | --- | --- | --- |
| 5X under-saturated | Species 1: **9.8** (9.76, 9.9)<br>Species 2: **10.5** (10.4, 10.6) | Species 1: **1.8** (1.77,1.8)<br>Species 2: **2.0** (1.97,2.1) | Species 1: **222** (217,227)<br>Species 2: **293** (280,306) | Species 1: **205** ± 16<br>Species 2: **239** ± 28 | Species 1: **204**<br>Species 2: **228** |
| 10X saturated | **12.08** (12.07, 12.9) | **1.73** (1.72,1.75) | **335** (332,306) | **352** ± 25 | **350** |
| 15X over-saturated | **13.76** (13.4, 14.1) | **1.63** (1.31,1.94) | **402** (287,517) | **365** ± 50 | **350** |
| 20X over-saturated | **14.3** (constant) | **1.65** (1.54,1.66) | **451** (447,455) | **369** ± 44 | **374** |

The saturated complex (**Figure 2, 10X TFAM**) displayed a remarkably homogenous s-value distribution, indicated by the vertical line in the van Holde Weischet plot, with 94% of the population sedimenting at 12 S, and a frictional coefficient of 1.7 (**Figure 2**, **Table 1**). In fact, the 95% confidence interval for the saturated complex (**Table 1**) is similar to that of eukaryotic nucleosomes which are known to behave as stable, uniform complexes^24,25^. An f/f0 of 1.7 suggests significant compaction of the DNA by TFAM (the f/f0 of free 207 bp DNA is ∼3.6, and an f/f0 of 1 indicates a perfect sphere with minimal viscous drag^24,25^). The calculated MW from SV-AUC (335 kDa) is also in close agreement with the MW of the saturated TFAM-160mer complex determined by mass photometry (352 kDa) (**Figure 1B**, **Figure 2**, **Table 1**). Together, these data confirm that 10 TFAM molecules compact the 160 bp DNA fragment into a complex that is homogenous in both size and shape.

As expected, the undersaturated complex (**Figure 2**, 5X TFAM) had a lower s-value and a higher f/f0, indicative of less compact species. Two main species were detected with calculated molecular weights that are consistent with mass photometry data at equivalent ratios (**Table 1**). Upon further TFAM titration beyond the saturation point (**Figure 2**, 15X and 20X TFAM), the s-value continued to increase without a significant increase in f/f0. However, in mass photometry experiments, oversaturating the complex did not increase the molecular weight (**Figure S2B** 15X and 25X, **Figure 2**, **Table 1**). This discrepancy is likely because SV-AUC experiments are carried out at a ∼60-fold higher concentration than mass photometry. The increase in molecular weight observed in SV-AUC is probably due to lower-affinity interactions that were not detectable by mass photometry. However, the f/f0 values indicate that excess TFAM binding did not further compact the DNA (**Table 1**). We suspect this is because these binding events at high concentrations and excess of TFAM are nonspecific and probably not physiologically relevant.

### Binding pattern and compaction are not sequence specific

Thus far, all DNA fragments used in this study were different lengths of the same randomized DNA sequence. Previous studies have shown that TFAM behaves differently on promoter sequences compared to nonspecific DNA sequences^20^. However, not all studies reported these differences and most were conducted on short DNA fragments that can accommodate 2 TFAM molecules at most^12,14,20^. To ensure that the uniform complexes we observed were not caused by sequence bias, we performed EMSAs and mass photometry with 160 bp DNA fragments derived from the mitochondrial genome (**Figure S3**). The ‘random position’ mtDNA fragment (“mtNS”) contains a region of the ND4 gene to represent an “average” stretch of DNA that TFAM would have to associate with to compact the genome. We also selected a 160 bp fragment from the D-loop region containing both the LSP and HSP promoters (“mtpromoter”) that are regulated by TFAM in its capacity as a transcription factor^1,9,10^.

EMSAs comparing the interaction of TFAM with random, mtNS and mtpromoter DNAs showed an identical binding pattern (**Figure S3A**). EMSA samples were also analyzed by mass photometry and the apparent molecular weights at each titration point were very similar between the three sequences (**Figure S2B, S4A,B**). The saturated complex on all three sequences showed a singular peak in mass photometry, indicative of a highly homogenous sample (**Figure S3B**). Prior studies observed a sequence bias in 1:1 complexes when TFAM oligomerization is not a factor^20^. Our data utilizing longer DNA demonstrates that TFAM does not have any sequence bias in how it oligomerizes on longer DNA, or in its ability to form homogenous complexes, in agreement with its role as a nucleoid compaction factor. As such, TFAM interaction with promoter DNA appears to be no different from its interaction with generic DNA.

### Micrococcal nuclease protection pattern confirms that TFAM forms higher order structures on DNA

We performed micrococcal nuclease (MNase) protection analysis of TFAM-160mer complexes to assess the DNA footprint protected by TFAM. Equilibrated complexes were subject to a titration of MNase followed by digestion of the protein and purification of the protected DNA fragments. The protected fragments were quantified on an Agilent TapeStation instrument which provides information on the length and relative abundance of each fragment. Three TFAM-160mer complexes at varying degrees of saturation were tested in this experiment: a half-saturated complex (4-5 TFAMs bound), a nearly-saturated complex (8 TFAMs bound) and a saturated complex (10 TFAMs bound). All three complexes yielded a regular digestion pattern showing the same protected fragments, indicating a regular TFAM distribution on the DNA (**Figure 3A**). As expected, increasing saturation resulted in increased protection of the full 160 bp fragment (**Figure 3B**). In the saturated complex, the 160 bp fragment was protected even at the highest MNase concentration, consistent with published data showing that fully compacted nucleoids are inaccessible^18^ (**Figure 3A-B**).

**Figure 3.**
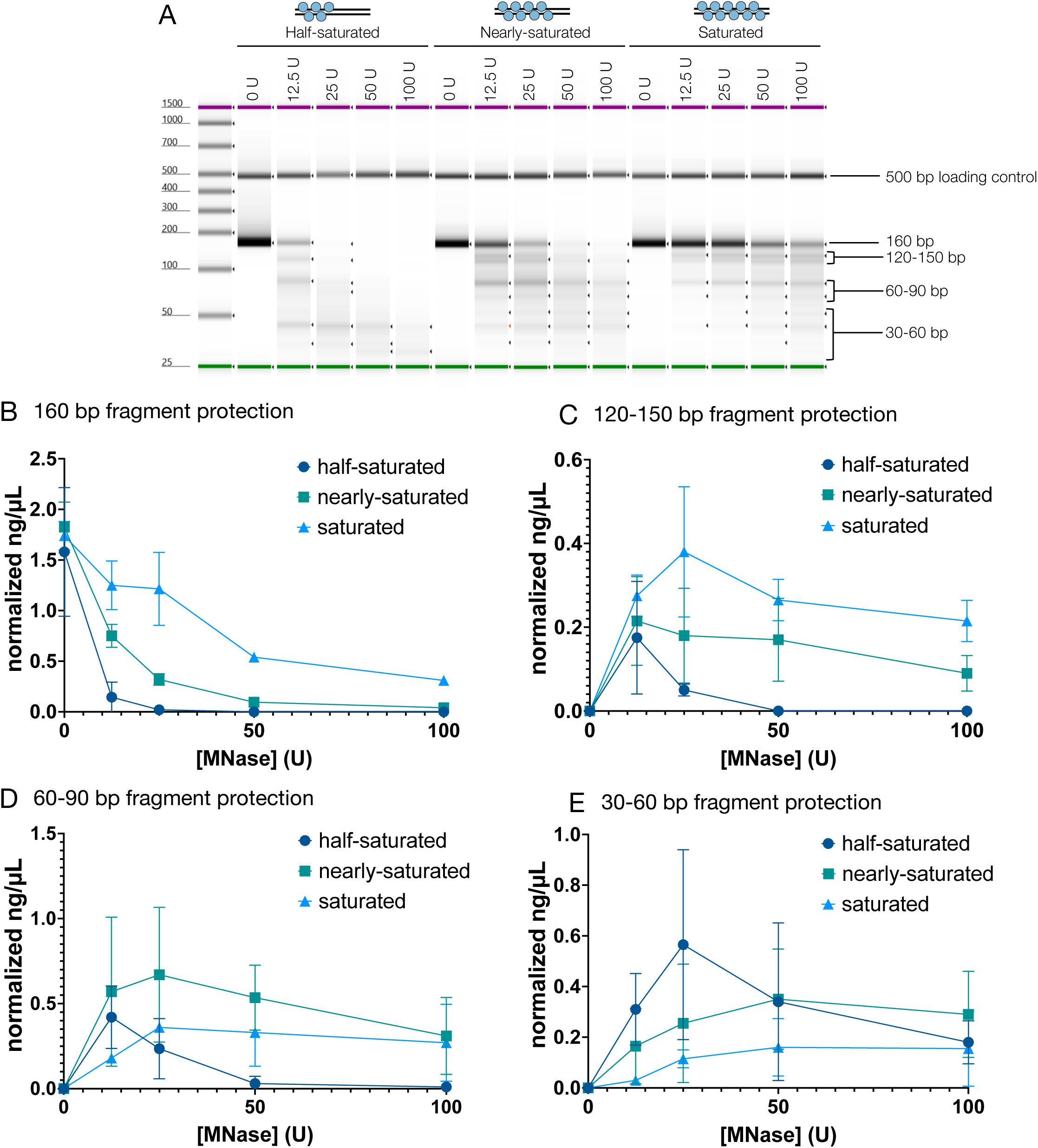
MNase digest analysis shows DNA protection in higher order structures. **A.** A representative capillary gel from the Agilent TapeStation showing the digest products. Cartoon above each complex depicts the expected number of TFAMs bound in each complex. Number of TFAMs expected was obtained from mass photometry experiments at the same TFAM:DNA ratio. Bands on the gel corresponding to the bins used in the analysis are highlighted. Most prominent band in the 60-90 bp bin range is ∼85 bp, and the most prominent band in the 30-60 bp bin is ∼45 bp. **B-E.** Normalized concentration for the indicated fragment at each TFAM:DNA ratio. Concentration is normalized to the loading control. n=2.

It is assumed that the nucleoid is compacted by a TFAM monomer inducing U-turns in 22-28 bp stretches of DNA^11–15^. In agreement with this, a 30 bp protected fragment was observed in all complexes tested in this experiment, although these were not as prominent as the larger fragments (**Figure 3A, 3E**). However, based on the 15 bp/TFAM footprint calculated from two orthogonal approaches, each 30 bp fragment would have 2 TFAMs bound. We also observed prominent protected fragments of ∼85 bp and ∼130 bp in length that were higher in abundance in the saturated and near-saturated complexes, consistent with the formation of higher order species that protect DNA (**Figure 3B-D**).

### Structural characterization of TFAM-DNA complexes reveals conformational heterogeneity

We utilized cryo-EM to assess the structure of the saturated TFAM-160mer complex. The native complex was unstable on cryo-EM grids, likely due to denaturation at the air water interface. As such, we performed Gradient Fixation (GraFix) with glutaraldehyde to visualize the crosslinked complex on regular EM grids (**Figure S5)**. We also utilized streptavidin monolayer affinity grids together with biotinylated TFAM to visualize the native complex^26^ (**Figure S6**).

Analysis of the crosslinked sample data yielded a featureless map showing only the overall shape of a complex (**Figure 4A**). Refinement of this density revealed high B-factors (> 500 Å^2^), indicative of a low confidence map (**Figure S5D**). The native complex, immobilized on streptavidin grids, yielded only fragmented maps showing the same overall size of a complex (**Figure 4B**). Similar to the data obtained from the crosslinked complex, refinement yielded maps with high B-factors (**Figure S6**). We also attempted data processing in Relion^27^ to validate that the poor map quality was not due to issues with particle alignment, but no improvement in map quality was observed. We also tested many other grid conditions with both native and crosslinked complexes, various processing strategies, as well as other DNA lengths and sequences, however no higher resolution map could be obtained.

**Figure 4.**
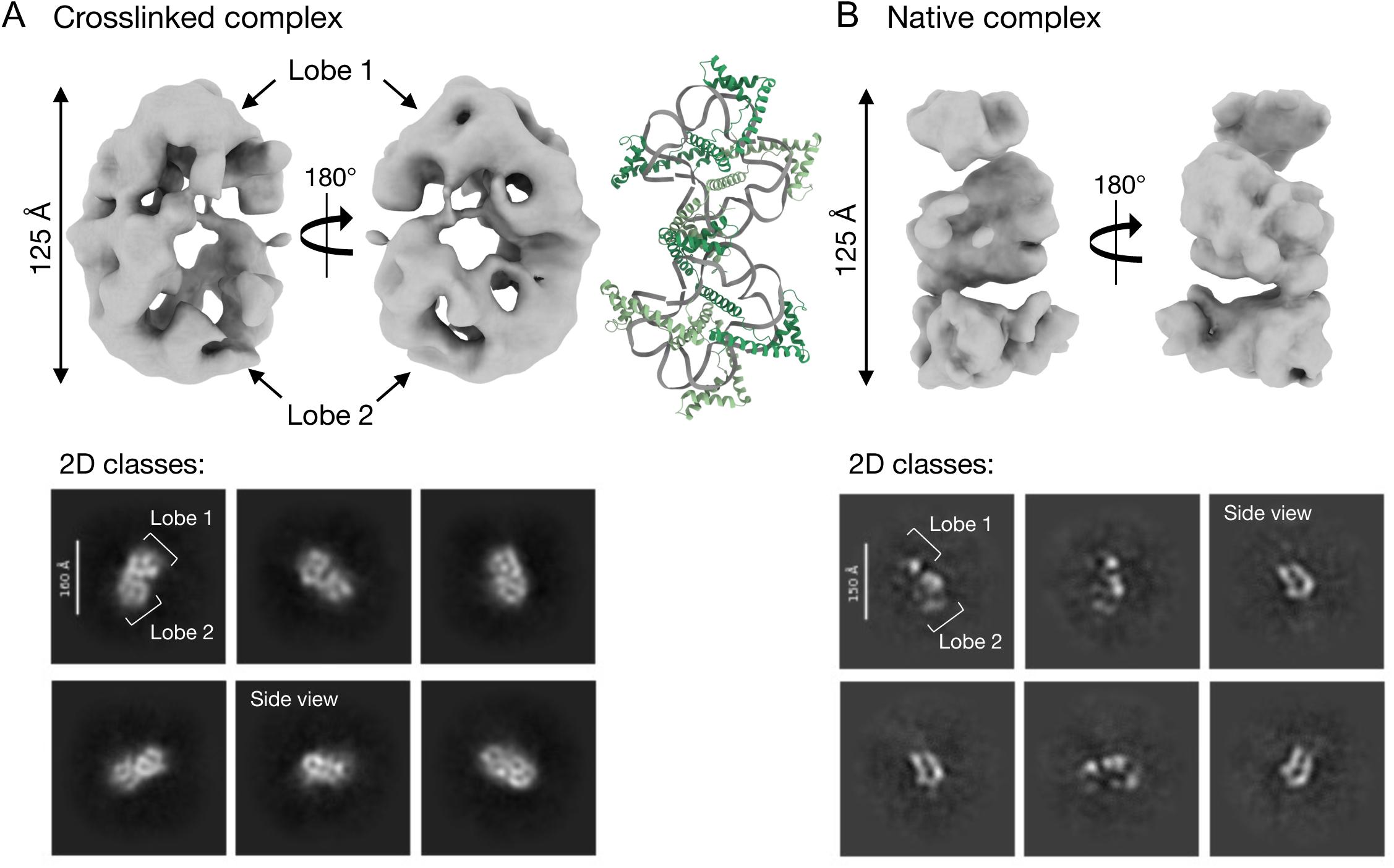
cryo EM analysis of crosslinked and native TFAM-160mer complexes. **A.** (Top) Electron density map of the crosslinked TFAM-160mer complex from an *ab initio* reconstruction from Cryosparc (133,739 particles). (Bottom) 2D classes of the same complex, obtained from cryosparc. Lobes and side views are indicated. **B.** (Top) Electron density map of the native TFAM-160mer complex from an *ab initio* reconstruction from Cryosparc (164,572 particles). (Bottom) 2D classes of the same complex from cryosparc. Putative lobes and side views are indicated. A TFAM-DNA crystal structure (PDB ID: 4NOD) is shown in the middle for size comparison. Note that this structure comprises four 22 bp DNA fragments with four TFAM molecules, forming two tight circles held together by TFAM dimerization in the crystal lattice.

Despite the lack of structural information from these maps, some information regarding the shape of the overall complex is discernable from 2D classifications from both datasets. 2D classes revealed a bi-lobed dumbbell-like complex (**Figure 4**). Potential side views of the complex were also visible in the 2D classes. Additionally, two lobes were visible in the reconstructed volume of the crosslinked complex (**Figure 4A**).

Although we cannot trace the trajectory of the DNA in our structures, it is apparent that it is highly bent and distorted in our densities, similar to what is observed in crystal structures of a single TFAM molecule on short DNA (**Figure 4, see discussion**).

Given the striking homogeneity of our saturated complexes observed in biochemical assays, we hypothesized that our inability to obtain good consensus electron density maps was due to inherent conformational heterogeneity in the complex. To test this hypothesis, we analyzed maps from both data sets in cryoDRGN^28^ (**Figure S7**). Most tools that model heterogeneity (for example, 3D variability analysis in cryosparc^29^) require particle orientations to be determined *a priori* and require the user to input a number of expected classes. CryoDRGN on the other hand reconstructs 3D volumes and learns a neural model of the distribution of 3D structures present in the data without requiring a number of expected classes. In this manner, cryoDRGN can model both discrete and continuous conformational heterogeneity. If discrete conformational states are present, these would appear as separate clusters in the latent space representation (the low dimensional map encoding the conformation of each particle), and these could then be determined as separate models. For both the crosslinked and native datasets, the distribution of particles in latent space is smooth with no distinct classes (**Figure S7**), suggesting continuous heterogeneity of the complex.

## Discussion

TFAM-mediated packaging of the mitochondrial genome into nucleoids is vital for the maintenance of mtDNA and consequently crucial for sustaining cellular energy production^1^. Prior biophysical analyses have focused on 1:1 TFAM-DNA complexes with short DNA fragments^11–15,20^. However, it is estimated that in the cell ∼1000 TFAM molecules are bound to a single copy of the 16.5 kb mtDNA in fully compacted nucleoids^6^. In this study, we investigate TFAM complexes with DNA fragments that are long enough to bind multiple TFAM molecules, to understand how it might organize DNA.

Our biochemical analysis demonstrates that TFAM cooperatively oligomerizes on the DNA to form complexes that are strikingly homogenous and compact in solution. Using mass photometry, we demonstrate that TFAM forms homogenous complexes on DNA fragments of different lengths (**Figure 1B**). The footprint of TFAM in the saturated complexes for all DNAs is consistently ∼15 bp/TFAM. This footprint differs from the 22-28 bp footprint observed in all published 1:1 crystal structures, suggesting that TFAM adopts a more crowded binding mode when oligomerizing on DNA. It is also worth noting that a footprint of ∼15 bp/TFAM corresponds to ∼1100 TFAM molecules per mtDNA which is consistent with published calculations of 1000-1500 TFAM molecules per mtDNA^6^.

Using AUC, we confirm the stoichiometries obtained by mass photometry and also show that TFAM-DNA complexes form a homogenously compact shape based on frictional ratio (f/f0) values (**Figure 2**). The confidence intervals for both sedimentation coefficient and f/f0 were similar to values obtained for nucleosomes and we thus deem these results derived from first principle very reliable^24,25^. Together, mass photometry and AUC data contradict the prevailing assumption that TFAM forms heterogenous complexes on DNA, suggesting that there may be a more regular organization to mitochondrial nucleoids than previously thought.

EMSA and mass photometry analysis of TFAM-DNA complexes comparing DNAs containing promoter sequences to nonspecific sequences showed that TFAM oligomerization does not appear to have any sequence specificity (**Figure S3**). This observation is in line with TFAM’s ability to coat the entirety of mtDNA regardless of sequence, as befits a nucleoid-associated protein (NAP), and consistent with previous in vitro studies showing that DNA bending is a general feature of TFAM independent of DNA sequence^12^. Yet, TFAM must somehow retain its specificity for promoter regions when functioning as a transcription factor. How TFAM switches between these two functions and binding modes is still unknown.

Using MNase protection assays, we show that TFAM forms higher order complexes that protect DNA against digestion (**Figure 3**). Similar MNase digest experiments in prior studies show some evidence of higher order species formation, but these species did not persist at later time points/higher enzyme concentrations^23^. In contrast, in our experiments, the larger protected fragments (>30 bp) persist even at the highest MNase concentration in saturated TFAM-DNA complexes.

Finally, our cryo-EM analysis of both native and crosslinked complexes provides evidence that TFAM exhibits conformational heterogeneity (**Figure 4, Figure S7**). Prior investigation of TFAM dynamics from single molecule FRET studies, using 1:1 complexes with short DNAs suggested two discrete conformational states^20^, however we do not observe discrete states here. Rather, we propose that

TFAM is continuously fast-exchanging and samples different conformations on the DNA, resulting in blurred, poor quality electron density map reconstructions (**Figure 4**). This appears to be an inherent feature of TFAM-DNA complexes, as they could not be resolved despite the significant effort invested into grid optimization. We assume that homogenous complexes were still detected in mass photometry and AUC because the continuous exchange of TFAM does not alter the overall dimensions and compaction of the complex.

A long-standing question in the field is whether there is any repeating organization in how TFAM compacts mtDNA, similar to how histones compact nuclear DNA into structural units of nucleosomes. We observe strong protection of 85 bp and 130 bp fragments in MNase digests (**Figure 3**). Analysis of the sedimentation behavior by AUC of equivalent complexes indicates that TFAM organizes the full length of the 160 bp DNA into a compact molecule (**Figure 2**) that is likely comprised of two of these ∼80 bp units observed in cryo-EM, with each containing 5 TFAM molecules. The two lobes in our 2D classes and 3D reconstructions appeared to be of similar size, however the exact length of DNA occupied in each of these lobes cannot be confidently discerned at the current resolution. The structure most consistent in shape with our densities comprises four 22 bp DNA fragments in complex with 4 TFAM molecules (PDB ID: 4NOD). In this crystal structure, the DNA forms two ‘closed circles’, held together by TFAM dimerization^12^. Dimerization was shown to be essential for DNA compaction using Tethered Particle Motion (TPM). While the path of the DNA is necessarily different in our structure because a continuous 160 bp DNA fragment was used, these similarities lead us to conclude that our densities likely only accommodate ∼80 bp of highly bent DNA, corresponding to ∼50% of the expected complex (**Figure 4**). We speculate that each lobe contains ∼40 bp of DNA and based on our calculated footprint (15 bp/TFAM), we expect 5 TFAMs bound in our density. Taking published structures into account, it is likely that TFAM dimerization contributes to DNA compaction.

In conclusion, our study demonstrates that TFAM oligomerizes on longer DNA into compact complexes that are strikingly homogenous in overall shape and size yet exhibit significant conformational heterogeneity that precludes structure determination by electron microscopy. This heterogeneity might also explain why no crystal structures of TFAM bound to longer DNA have been published. While a high-resolution structure may not be possible using cryo-EM due to these dynamics, future studies should utilize methods such as NMR spectroscopy to gain insight into the three-dimensional arrangement and binding surfaces of TFAM in these complexes.

## Methods

### TFAM expression & purification

TFAM lacking the mitochondrial targeting sequence (residues 43-246) was cloned into a pET expression vector through restriction-free cloning. The protein was expressed in 1 L cultures of BL21(DE3) *E.coli* cells grown from overnight cultures. Cells were induced with 0.2 mM IPTG at OD ∼1.0. Cells were collected after 3 hours by centrifugation at 6000 rpm for 30 min at 4 °C. Media was decanted and the pelleted cells were flash frozen in liquid nitrogen and stored at −80 °C until thawed for purification.

Cells were thawed on ice and resuspended in lysis buffer (20 mM HEPES pH 7.5, 50 mM NaCl, 1 mM EDTA and 1 Pierce Complete protease inhibitor tablet per 50 ml). Cells were then lysed by sonication (3 rounds of 1s on/off for 1 min). The lysate was centrifuged at 16000 rpm for 30 min at 4 °C and the resulting supernatant was filtered through a 0.45 µm syringe filter.

All column purification steps were performed on an AKTA Pure FPLC. Filtered lysate was applied to a 5 mL SP HP column (Cytiva) in buffer A (20 mM HEPES pH 7.5, 50 mM NaCl, 1 mM EDTA, 1 mM TCEP and 1 mM AEBSF) and eluted using a linear gradient of buffer B (20 mM HEPES pH 7.5, 1 M NaCl, 1 mM EDTA, 1 mM TCEP and 1 mM AEBSF). Fractions were analyzed by SDS PAGE, appropriate fractions were pooled, diluted in buffer A to ∼50-100 mM NaCl and applied to a 5 mL HiTrap Heparin HP column (Cytiva). Protein was eluted using a linear gradient ranging of buffer B (same buffers as SP column). Fractions were analyzed by SDS PAGE, appropriate fractions were pooled and concentrated to ∼1 mL. Concentrated fractions were loaded onto a 120 mL S200 column (Cytiva) in storage buffer (20 mM HEPES pH 7.5, 300 mM NaCl, 1 mM TCEP and 1 mM AEBSF). Fractions were analyzed by SDS PAGE. Appropriate fractions were pooled and concentrated to a desirable working concentration. Concentrated protein was aliquoted, flash frozen in liquid nitrogen and stored at −80 °C.

### DNA amplification and purification

#### Randomized sequence DNA fragments

A 500 bp DNA fragment of a randomized sequence was computationally generated. In this sequence, each 10 bp stretch has a GC content of 50%. One forward primer and 26 different reverse primers corresponding to different lengths of DNA were used to generate a “random DNA” library consisting of 26 different lengths. 80 bp, 160 bp, 240 bp and 400 bp fragments from this library were used in this study. For fluorescent DNAs, the forward primer was labeled with Alexa 488.

DNAs were amplified by PCR using the OneTaq Master Mix (New England Biolabs). Reactions were pooled and applied to an 8 mL Mono Q column (Cytiva) in wash buffer (10 mM HEPES pH 7.5, 1 mM EDTA). DNA was eluted using a linear gradient of elution buffer (10 mM HEPES pH 7.5, 1 M NaCl, 1 mM EDTA). Fractions were analyzed on 10% Native TBE gels, pooled and further purified by ethanol precipitation.

<u>80mer:</u> 5’-GCTAGTCCGTCTTCTACTCTGAAATGAGCAGTCCTAGTCAGCAAGATCGCTCAGCCAACT TTCTACCAGCGCAACCCTAA-3’

<u>160mer:</u> 5’-GCTAGTCCGTCTTCTACTCTGAAATGAGCAGTCCTAGTCAGCAAGATCGCTCAGCCAACT TTCTACCAGCGCAACCCTAATCTACCCCATGAATGAAGCCGCACCCAAAACCGCATTCTAAG GAGTGACATTAACCCTCGGTGAGGATGTCCATACAAGC-3’

<u>240mer:</u> 5’-GCTAGTCCGTCTTCTACTCTGAAATGAGCAGTCCTAGTCAGCAAGATCGCTCAGCCAACT TTCTACCAGCGCAACCCTAATCTACCCCATGAATGAAGCCGCACCCAAAACCGCATTCTAAG GAGTGACATTAACCCTCGGTGAGGATGTCCATACAAGCACCTCCTACTACGGATCGAACCGT TAGTTCCCCAACTAAGTCCAAACCGTTAGACCGCTTTCCGTACCATTCCGGTACTT-3’

<u>400mer:</u> 5’-GCTAGTCCGTCTTCTACTCTGAAATGAGCAGTCCTAGTCAGCAAGATCGCTCAGCCAACT TTCTACCAGCGCAACCCTAATCTACCCCATGAATGAAGCCGCACCCAAAACCGCATTCTAAG GAGTGACATTAACCCTCGGTGAGGATGTCCATACAAGCACCTCCTACTACGGATCGAACCGT TAGTTCCCCAACTAAGTCCAAACCGTTAGACCGCTTTCCGTACCATTCCGGTACTTATCTTCG CCACAACCTGAGACAATCCCAAGCTTAAGGCTCGACACAGACTGACGAAGGATATATCTCGC CCTAACCGTACCTCTATACCGCCATGAAGGAAGTGCCAAGTAGCCACAGAACCTTGGGATAG CAAGACTCTATGTCCCAGACCTCACTAAC-3’

#### Mitochondrial DNA-derived sequence DNA fragments

For the “mtpromoter” sequence, a 400 bp gBlock containing both promoter sequences (HSP and LSP) was designed along with primers such that either a 160 bp or a 400 bp DNA fragment containing both promoters could be amplified. For the “mtNS” sequence, gBlock containing 400 bp of the ND4 gene was designed along with primers such that either a 160 bp or a 400 bp DNA fragment could be amplified. For fluorescent DNAs, the forward primer was labeled with Alexa 488.

All DNAs were amplified by PCR using the OneTaq Master Mix (New England Biolabs). Reactions were pooled and applied to an 8 mL Mono Q column (Cytiva) in wash buffer (10 mM HEPES pH 7.5, 1 mM EDTA). DNA was eluted using a linear gradient of elution buffer (10 mM HEPES pH 7.5, 1 M NaCl, 1 mM EDTA). Fractions were analyzed on 10% Native TBE gels, pooled and further purified by ethanol precipitation.

<u>mtpromoter 160mer:</u> 5’-TTTGGCGGTATGCACTTTTAACAGTCACCCCCCAACTAACACATTATTTTCCC CTCCCACTCCCATACTACTAATCTCATCAATACAACCCCCGCCCATCCTACCCAGCACACACA CACCGCTGCTAACCCCATACCCCGAACCAACCAAACCCCAAAGA-3’

<u>mtpromoter 400mer:</u> 5’-CAAAAACAAAGAACCCTAACACCAGCCTAACCAGATTTCAAATTTTATCTTT TGGCGGTATGCACTTTTAACAGTCACCCCCCAACTAACACATTATTTTCCCCTCCCACTCCCA TACTACTAATCTCATCAATACAACCCCCGCCCATCCTACCCAGCACACACACACCGCTGCTAA CCCCATACCCCGAACCAACCAAACCCCAAAGACACCCCCCACAGTTTATGTAGCTTACCTCC TCAAAGCAATACACTGAAAATGTTTAGACGGGCTCACATCACCCCATAAACAAATAGGTTTG GTCCTAGCCTTTCTATTAGCTCTTAGTAAGATTACACATGCAAGCATCCCCGTTCCAGTGAGT TCACCCTCTAAATCACCACGATCAAAAGGAACAAG-3’

<u>mtNS 160mer:</u> 5’-ATGCTAAAACTAATCGTCCCAACAATTATATTACTACCACTGACATGACTTTCCAAA AAACACATAATTTGAATCAACACAACCACCCACAGCCTAATTATTAGCATCATCCCTCTACTAT TTTTTAACCAAATCAACAACAACCTATTTAGCTGTTCCC-3’

<u>mtNS 400mer:</u> 5’-ATGCTAAAACTAATCGTCCCAACAATTATATTACTACCACTGACATGACTTTCCAAA AAACACATAATTTGAATCAACACAACCACCCACAGCCTAATTATTAGCATCATCCCTCTACTAT TTTTTAACCAAATCAACAACAACCTATTTAGCTGTTCCCCAACCTTTTCCTCCGACCCCCTAA CAACCCCCCTCCTAATACTAACTACCTGACTCCTACCCCTCACAATCATGGCAAGCCAACGCC ACTTATCCAGTGAACCACTATCACGAAAAAAACTCTACCTCTCTATACTAATCTCCCTACAAA TCTCCTTAATTATAACATTCACAGCCACAGAACTAATCATATTTTATATCTTCTTCGAAACCAC ACTTATCCCCACCTTGGCTATCATCA-3’

### Fluorescence Polarization (FP)

TFAM was titrated into 10 nM of Alexa488 labeled DNA substrate in binding buffer (20 mM HEPES pH 7.5, 100 mM NaCl, 1 mM EDTA) and incubated at room temperature for 30 min. FP was measured using a BMG Labtech CLARIOstar microplate reader. Data were analyzed in GraphPad Prism (v10.4.2) and fit to a four-parameter Hill Equation

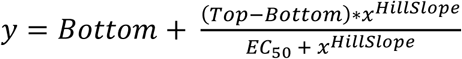

### Electro-Mobility Shift Assays (EMSA)

TFAM (25 µM or 50 µM, depending on DNA length) was titrated into 1 µM of DNA substrate in binding buffer (20 mM HEPES pH 7.5, 100 mM NaCl, 1 mM EDTA) at the indicated molar ratios and incubated at room temperature for 30 min. Samples were mixed 1:1 with 50% sucrose and loaded onto 6% DNA retardation gels (Invitrogen) and run at 100 V for 60 min in 0.2X TBE at room temperature. Gels were stained with Ethidium Bromide.

### Mass photometry

#### Sample preparation and data collection

Mass photometry measurements were performed on a Refeyn TwoMP mass photometer (Refeyn Ltd). Glass coverslips were cleaned with isopropanol and deionized water, then dried with N_2_ gas. Coverslips were coated with a 0.01% Poly-L-lysine solution (Sigma #P4707) for 25 seconds, rinsed with deionized water and dried with N_2_ gas.

Samples were prepared for EMSAs as described above. Samples were diluted to a final concentration of ∼5 nM onto the slide. All dilutions were performed at room temperature in binding buffer (20 mM HEPES pH 7.5, 100 mM NaCl). A 60 s video was recorded immediately after the final dilution, using Refeyn AcquireMP. ß-amylase was used as a mass standard to convert contrast signal to molecular weight. This was executed for each titration point for all four DNA lengths tested. Performed in triplicate for 80mer and 160mer; performed in duplicate for 240mer and 400mer.

#### Data analysis

Using the merge function in the Refeyn Discover MP software, we combined replicate data for each titration point prior to molecular weight calculation. The Gaussian function in the DiscoverMP software was used to calculate molecular weights from particle count vs molecular mass distribution histograms. The number of TFAMs bound in each complex was calculated by subtracting the theoretical molecular weight of the DNA and dividing the remainder by the molecular weight of a TFAM monomer (24.5 kDa). The footprint of each TFAM molecule (#bp/TFAM) was calculated by dividing the DNA length by the number of TFAMs bound in the complex. The saturated complex for each DNA were determined based on the maximum molecular weight obtained with further TFAM titration, or the maximum molecular weight obtained prior to a large increase in background and decrease in main peak signal.

### Sedimentation Velocity Analytical Ultracentrifugation (SV-AUC)

We performed SV-AUC with absorbance optics (λ = 260 nM). All samples were prepared at the indicated TFAM:DNA ratios at a final DNA concentration of 390 nM in binding buffer (20 mM HEPES pH 7.5, 100 mM NaCl, 1 mM EDTA). Samples were spun at 30,000 rpm at 20 °C in the Beckman XL-A ultracentrifuge using the An60Ti rotor in binding buffer. Data analysis was performed in Ultrascan III v4.0^30^ to determine sedimentation coefficients (S(20,W)), frictional ratios (f/f0) and molecular weights for each sample. Van Holde Weischet plot and distribution plots were generated in GraphPad Prism (v10.4.2).

### Micrococcal nuclear (MNase) digest protection assay

#### Sample preparation and digest reaction

Samples were prepared in binding buffer (20 mM HEPES pH 7.5, 100 mM NaCl, 1 mM EDTA) at a final DNA concentration of 1 µM at the indicated TFAM:DNA ratios and incubated at room temperature for 30 min prior to digest reaction assembly. MNase (New England Biolabs) was diluted to four working concentrations: 100 U/µL, 50 U/µL, 25 U/µL and 12.5 U/µL in MNase reaction buffer (50 mM Tris-HCl pH 7.9, 5 mM CaCl2).

Each 100 µL digest reaction was assembled with 30 µL of the complex, 5 µL BSA (Thermofisher Scientific) and 1 µL of the desired MNase concentration. The remaining volume was made up with reaction buffer. Reactions were incubated at 37 °C for 10 min and quenched with 0.1 M EDTA (final concentration). As an internal loading control for downstream TapeStation analysis, 35 ng of the 500 bp DNA loading control was added to each reaction after quenching. Next, 20 µg of Proteinase K (New England Biolabs) was added to each reaction to digest proteins in the complex. Reactions were incubated at 50 °C for 30 min. Resulting DNA fragments were purified using the Qiagen MinElute kit following instructions provided with the kit.

#### TapeStation data collection and analysis

All digested samples were analyzed on an Agilent TapeStation 2200 to determine lengths and relative quantities of protected fragments. Samples were diluted for TapeStation analysis following the manufacturer’s specifications. Samples were analyzed on a D1000 ScreenTape (Agilent).

For each sample, the detected fragments were binned by 30 bp. This length was chosen from the assumption that TFAM exists as a dimer in these complexes, with each monomer bound to 15 bp, as determined by mass photometry. The total DNA concentration (ng/µL) in each bin was then divided by the concentration of the 500 bp loading control fragment to calculate the normalized ng/µL of each length range. Normalized ng/µL vs [MNase] plots were generated in GraphPad Prism (v10.4.2) for each TFAM:DNA ratio tested.

### Sucrose Gradient Fixation (GraFix) crosslinking

A 10-30% (w/v) sucrose gradient was prepared in 13.2 mL polypropylene tubes (Beckman Coulter 331372). To assemble the gradient, 6 mL of top buffer (20 mM HEPES pH 7.5, 100 mM NaCl, 1 mM EDTA, 10% sucrose) was added to the tube, followed by 6 mL of the bottom buffer (20 mM HEPES pH 7.5, 100 mM NaCl, 1 mM EDTA, 30% sucrose, 0.25% glutaraldehyde). Bottom buffer was added slowly to displace the top buffer until the demarcation line reached halfway. A gradient maker (BioComp Gradient Master) was used to generate a continuous gradient. 200 µL of the TFAM-160mer complex (3 µM) was applied to the top of the gradient and spun at 30,000 rpm for 18 hrs at 4 °C. Fractionation was performed using with absorbance optics (λ = 260 nM and λ = 280 nM) using the same gradient maker. Fractions were analyzed using SDS PAGE and 6% DNA retardation gels (Invitrogen) (Figure S4). Appropriate fractions were dialyzed into binding buffer (20 mM HEPES, 100 mM NaCl, 1 mM EDTA) to remove sucrose. After dialysis, samples were concentrated to a desired stock concentration.

### Biotinylated TFAM preparation

TFAM was biotinylated using the EZ-link Maleimide-PEG2-Biotin reagent (Thermofisher Scientific). 5 mg/mL TFAM was incubated with 1 mM of the reagent at room temperature for 2 hours in binding buffer (20 mM HEPES pH 7.5, 100 mM NaCl, 1 mM EDTA). Excess reagent was removed using a spin column and biotinylated TFAM was concentrated to a desired working concentration. Successful biotinylation was validated using SDS PAGE where a clear shift was visible with labeled TFAM (Figure S5A). We also validated that biotinylated TFAM bound DNA with the same affinity as WT TFAM using Fluorescence Polarization (Figure S5B).

### Single particle cryo-EM grid preparation, data collection and processing

#### GraFix TFAM-160mer complex

C-flat 1.2/1.3 300 mesh grids were glow discharged at 40 mA for 30s using a Tergeo-EM plasma cleaner. 4 µL of 2.5 µM sample from GraFix was applied to the grid, blotted (2s, blot force 3) and plunge frozen in liquid ethane using the VitroBot Mark IV set to 100% humidity at 4 °C.

Grids were screened using a Glacios TEM and final collection was performed on a Titan Krios G3i with a Falcon 4 detector. 17,742 movies were collected at a magnification of 130,000x (0.97 Å/px) using a dose of 50 e^−^/Å^2^ using EPU (Thermofisher). The defocus range was −0.6 µm to −1.6 µm.

Preprocessing (motion correction, CTF correction and micrograph curation) was performed in Cryosparc (v4.7.1)^31^. Particles were picked using blob picker and particles were curated using iterative 2D classification in both Cryosparc and RELION (v5.0)^27^. *Ab initio* reconstructions were performed in cryosparc, followed by homogenous and non-uniform refinements. In an effort to obtain better density maps, 3D classification was also performed in RELION using a negative stain density as the initial model but no higher quality maps were obtained.

#### Native TFAM-160mer complex

Quantifoil Au 1.2/1.3 streptavidin monolayer affinity grids were made in-house using previously described procedures. The TFAM-160mer complex was assembled at 250 nM in binding buffer (20 mM HEPES pH 7.5, 100 mM NaCl, 1 mM EDTA). 4 µL of the complex was incubated on the streptavidin monolayer affinity grids for 10 min in a humidifying chamber. The grid was then washed 3x in 100 µL drops of binding buffer and the remaining buffer was wicked away using Whatman filter paper after the last wash step. 4 µL of binding buffer was added immediately to the grid after wicking. The grid was then transferred to a Leica GP2 set to 90% humidity at 4 °C for plunge freezing. Grids were blotted for 2s and plunge frozen into liquid ethane.

Grids were screened using a Glacios TEM and final collection was performed on a Titan Krios G3i with a Falcon 4 detector. 16,605 movies were collected at a magnification of 130,000x (0.91 Å/px) using a dose of 50 e^−^/Å^2^ using EPU (Thermofisher). The defocus range was −0.6 µm to −1.6 µm.

Motion correction was performed in Cryosparc (v4.7.1)^31^. Background lattice subtraction was performed using scripts from Cookis et al (2023)^26^. Subtracted micrographs were re-imported into Cryosparc for CTF correction and micrograph curation. Particles were picked using blob picker and particles were curated using iterative 2D classification in both Cryosparc and RELION (v5.0)^27^. *Ab initio* reconstructions were performed in cryosparc, followed by homogenous and non-uniform refinements. In an effort to obtain better density maps, 3D classification was also performed in RELION using a negative stain density as the initial model but no higher quality maps were obtained.

### Heterogeneity analysis of cryo-EM data with cryoDRGN

Due to the inability to reconstruct a high confidence map, we analyzed both clean particle stacks in cryoDRGN^28^ (v3.5.4). Particle stacks were downsampled to 128 pixels in cryoDRGN. We trained the cryoDRGN heterogenous *ab initio* model on these particles on a 10D latent variable model for 50 epochs. The encoder and decoder architectures were 256 x 3. This analysis yielded no distinct populations for either dataset. We also tested a 1D latent variable model with the same architecture for 50 epochs and a larger architecture (encoder 256 x 3 and decoder 512 x 5) with a 1D latent variable model for 100 epochs. Neither of these produced distinct populations in the UMAPs or PCA projections. Evaluation of the resulting densities revealed poor quality maps and no differences between the maps at different Z values. Given the homogeneity of this complex in biochemical analyses, we concluded that the lack of distinct populations and poor reconstructions in cryoDRGN were due to inherent conformational dynamics in the complex. In other studies utilizing this tool with higher resolution data, different conformations of the complex can be visualized along the trajectory of the latent space to discern different binding modes^28,32^. However, due to the low resolution maps obtained herein, such analysis could not be performed in this case.

## Author contributions

**SRW:** conceptualization; methodology; investigation; data curation; validation; formal analysis; visualization; writing (original draft, review, and editing)

**WT:** investigation; data curation; validation; writing (review and editing)

**KL:** conceptualization; funding acquisition; project administration; supervision; writing (original draft, review, and editing)

## Data sharing plan

All raw cryo-EM data will be deposited in EMPIAR upon publication, along with density maps. All original data files will be submitted in a repository upon publication

## Acknowledgements

Cryo-EM imaging was assisted by Dr. Erik Hartwick, and data collection was performed at the Biochemistry Krios Electron Microscopy Facility (BioKEM) at CU Boulder (RRID: SCR_019057). We acknowledge Dr. Annette Erbse and the Shared Instruments Pool (RRID: SCR_018986) of the Department of Biochemistry at the University of Colorado Boulder for providing access to the Avanti JXN 26 Super Speed Centrifuges and rotors, funded by NIH Grant (R24OD033699-01). Software used in this project was curated by SBGrid. This work utilized the Blanca condo computing resource at the University of Colorado Boulder. Blanca is jointly funded by computing users and the University of Colorado Boulder. Data storage supported by the University of Colorado Boulder PetaLibrary. We thank Dr. Shawn Laursen for the development of the 50% G/C 150 bp sequence DNA utilized in assays. We thank Dr. Vignesh Kasinath and Dr. Eva Nogales for providing streptavidin monolayer affinity grids. SRW was a trainee of the Signaling and Cellular Regulation T32 training program for a duration of this work (T32 GM142607, SRW). Funded by the Howard Hughes Medical Institute (WT, KL).

---

**Figure S1.**
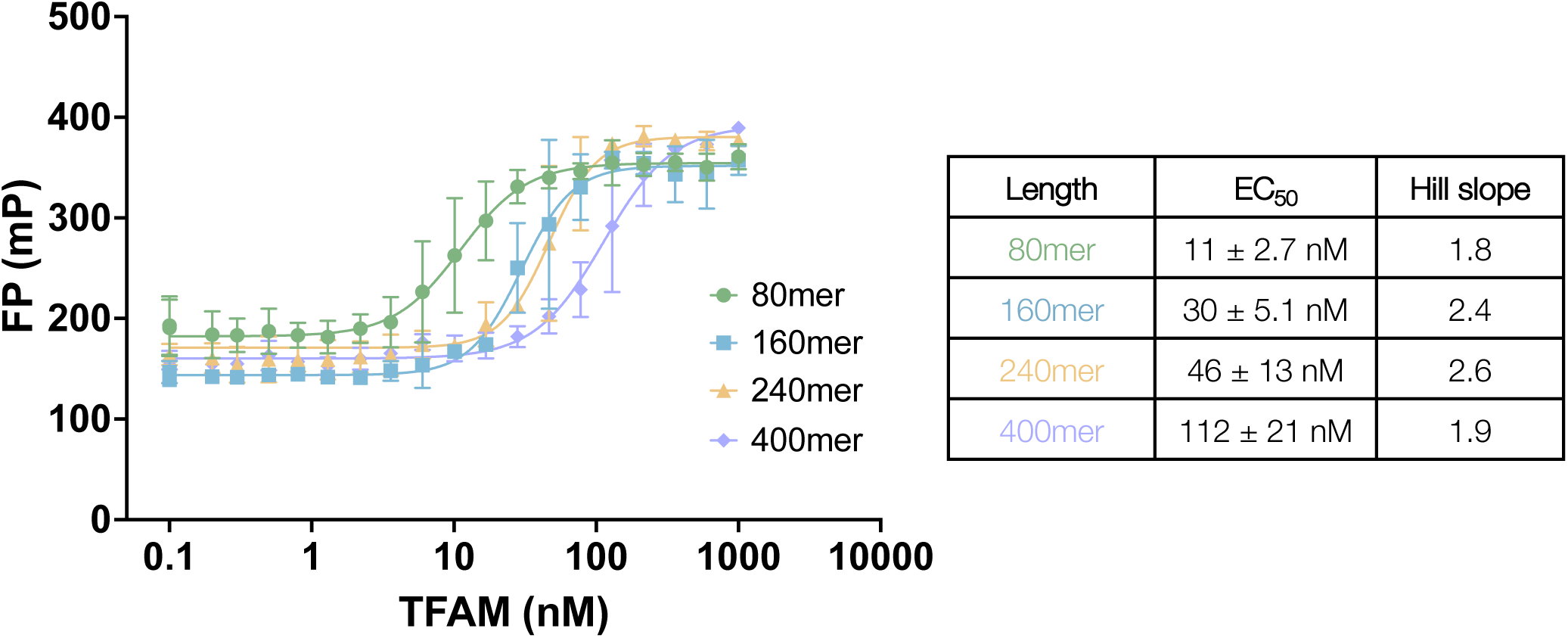
TFAM binds DNA cooperatively and with high affinity. Fluorescence Polarization (FP) analysis of TFAM with four DNA fragments of increasing length (randomized sequence). Data were fit to a four parameter Hill equation to obtain EC_50_ values and Hill slopes. n=2. Error shown in the inset was calculated from 95% confidence intervals.

**Figure S2.**
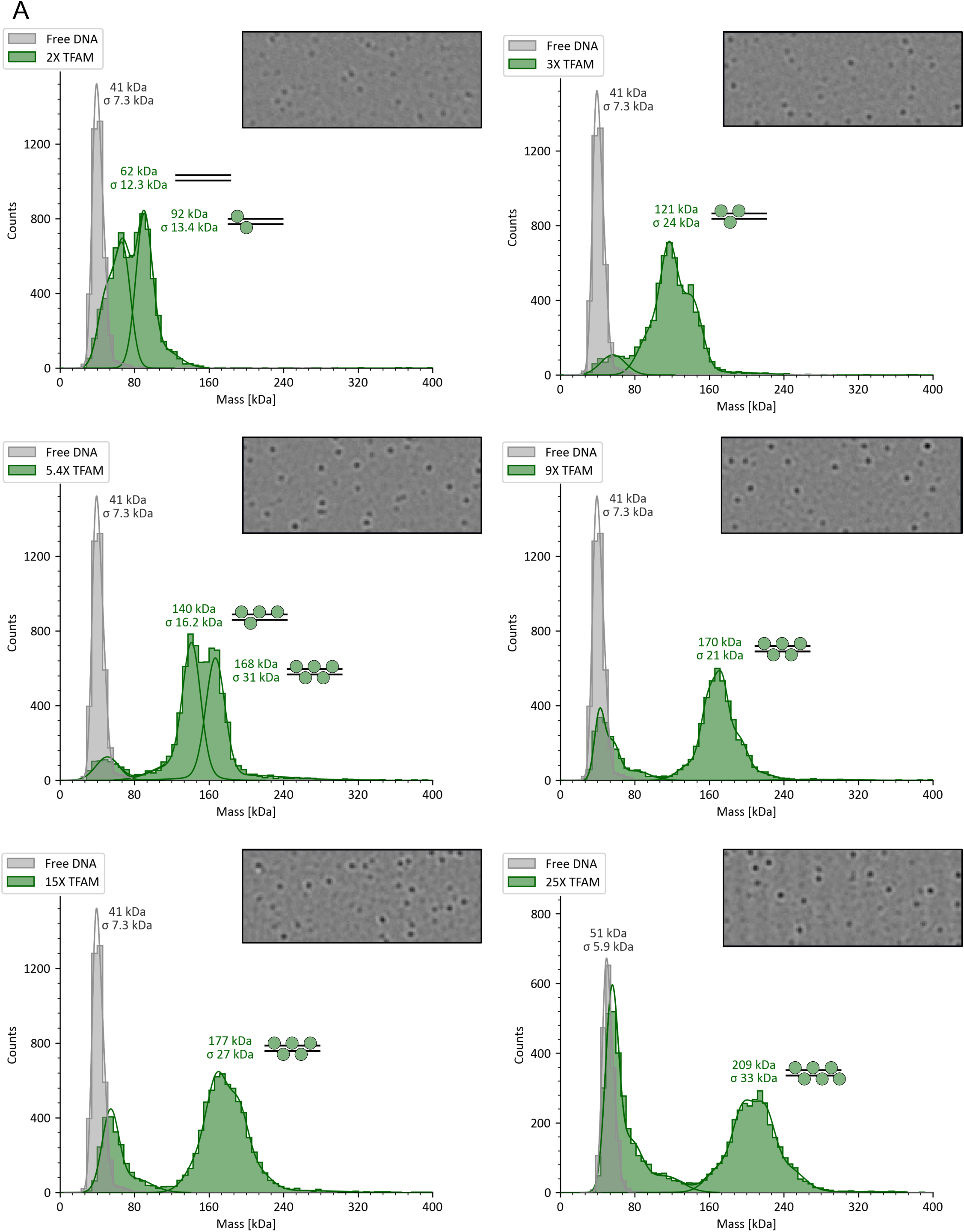
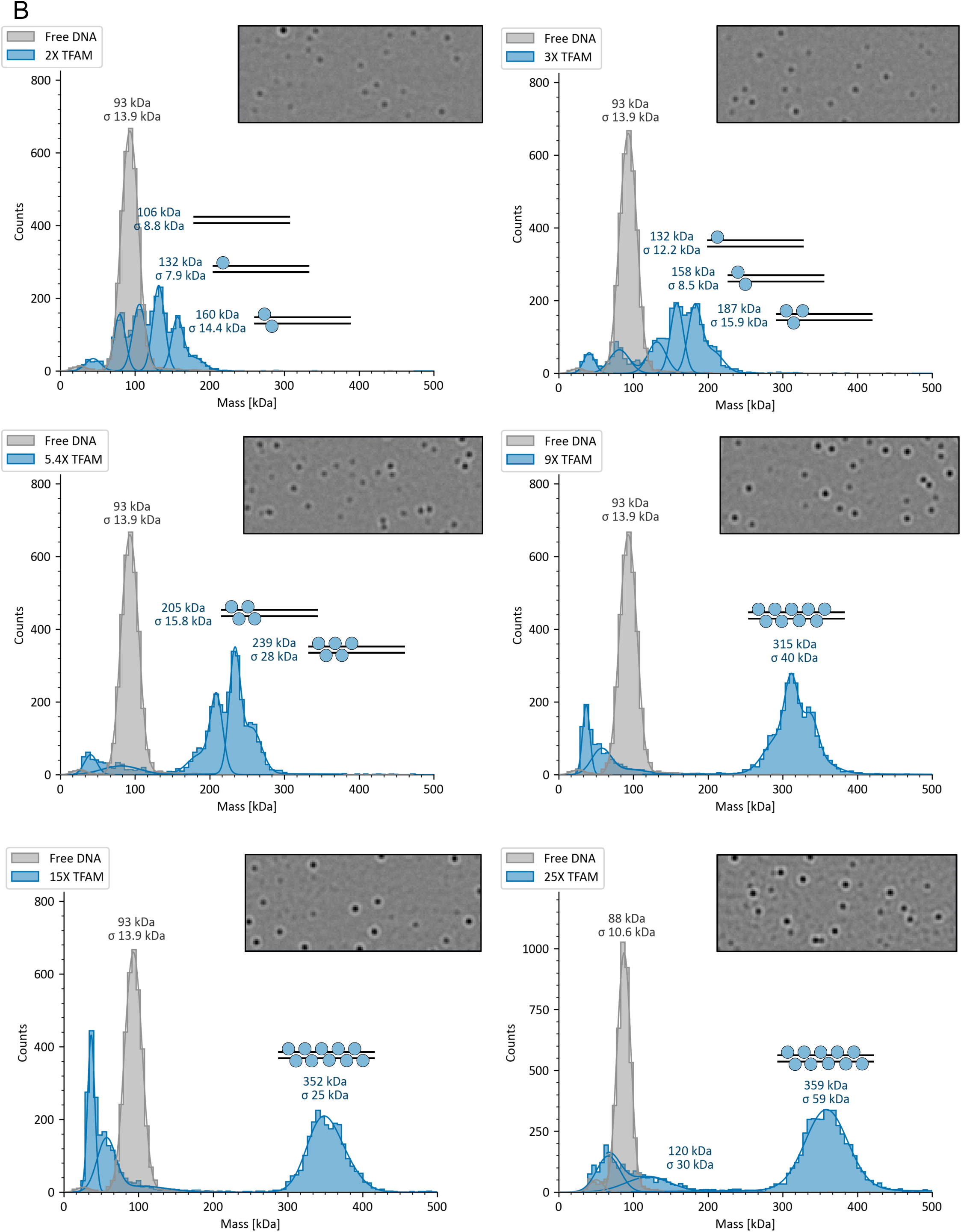
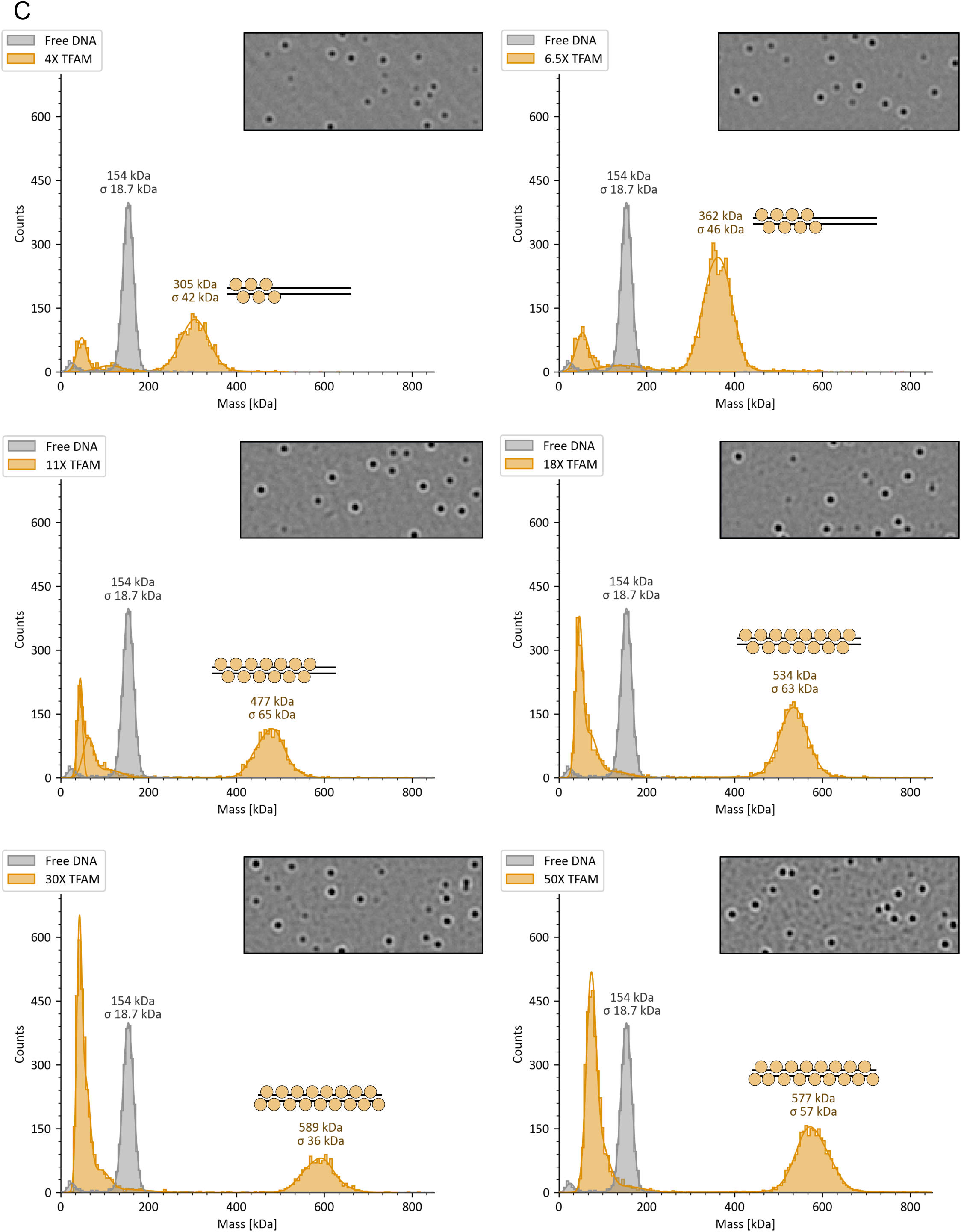
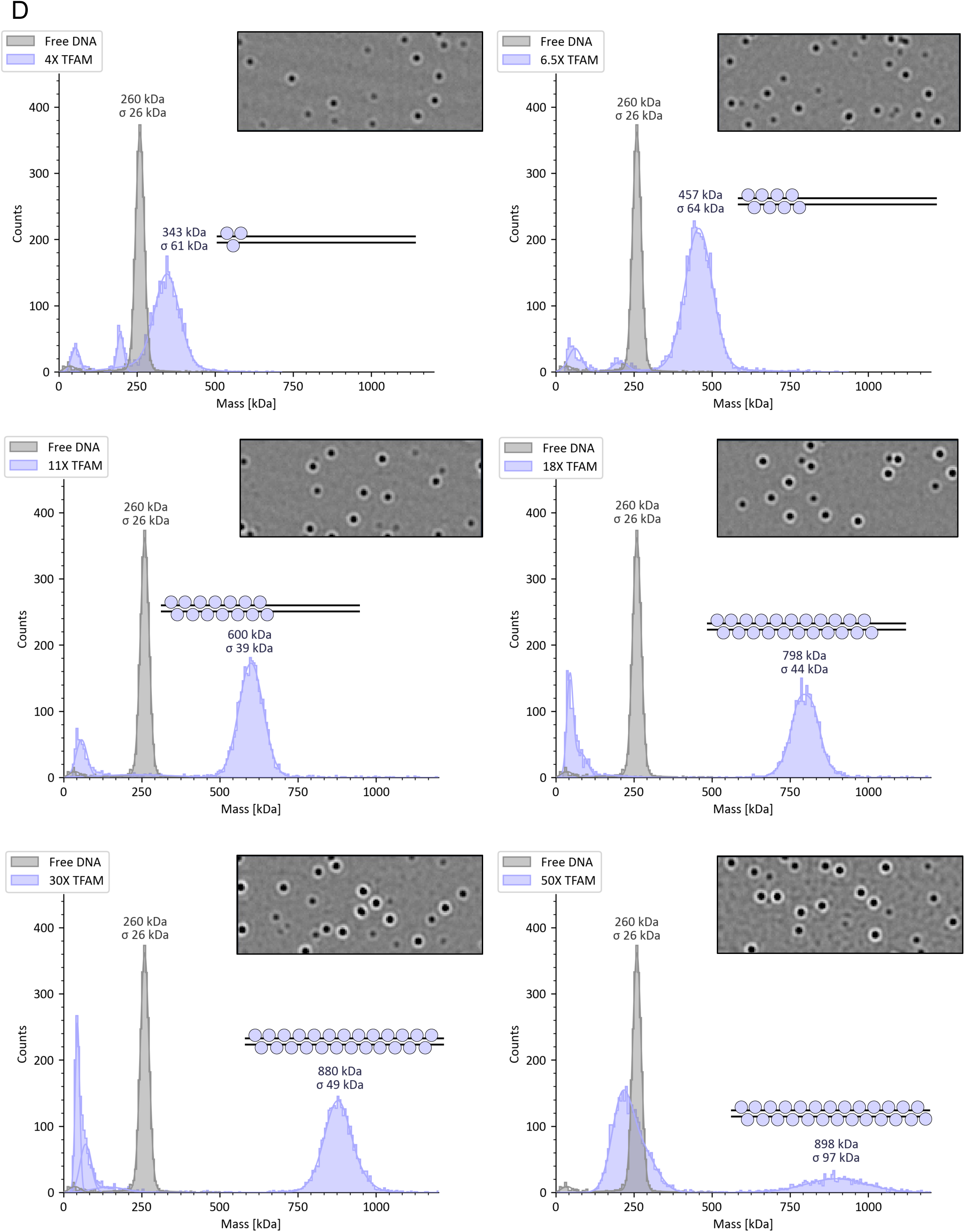
Mass photometry data for each titration point with DNA fragments of different lengths and with randomized sequences. **A.** TFAM titration into 80 bp DNA. **B.** TFAM titration into 160 bp DNA. **C.** TFAM titration into 240 bp DNA. **D.** TFAM titration into 400 bp DNA. Cartoon near each peak represents the number of TFAMs bound in each detected complex. Representative data shown.

**Figure S3.**
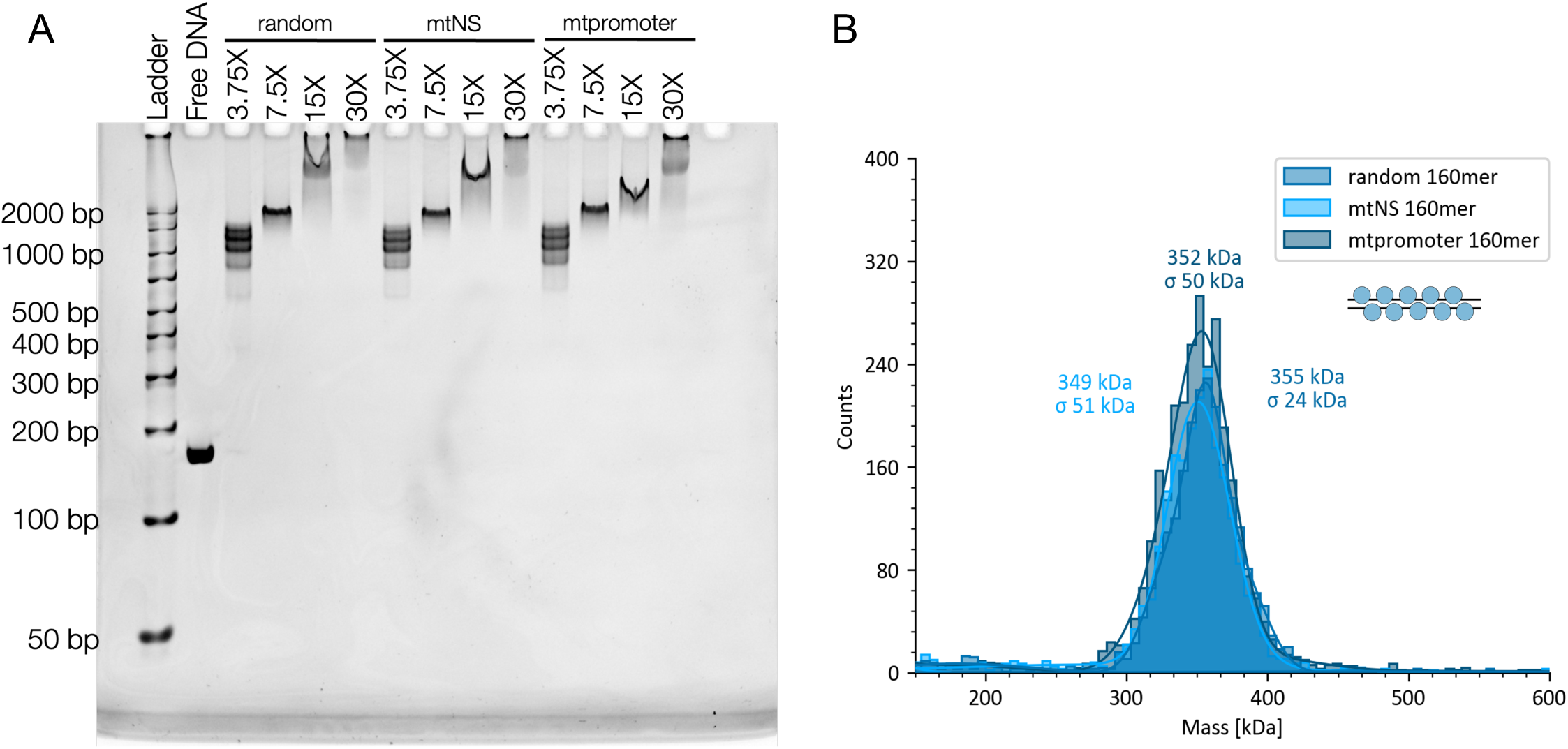
TFAM oligomerization is not sequence specific. **A.** EMSA with 160 bp fragments of different sequences: random DNA sequence, a nonspecific mtDNA sequence (“mtNS”), and a specific mtDNA sequence (“mtpromoter”). Samples were run on a 6% DNA retardation gel and visualized by ethidium bromide staining. Representative of n=2 shown. B. Mass photometry of the saturated complex (15X TFAM) with each DNA fragment. Representative of n=3 for random 160mer, and n=1 for mtNS and mtpromoter DNAs

**Figure S4.**
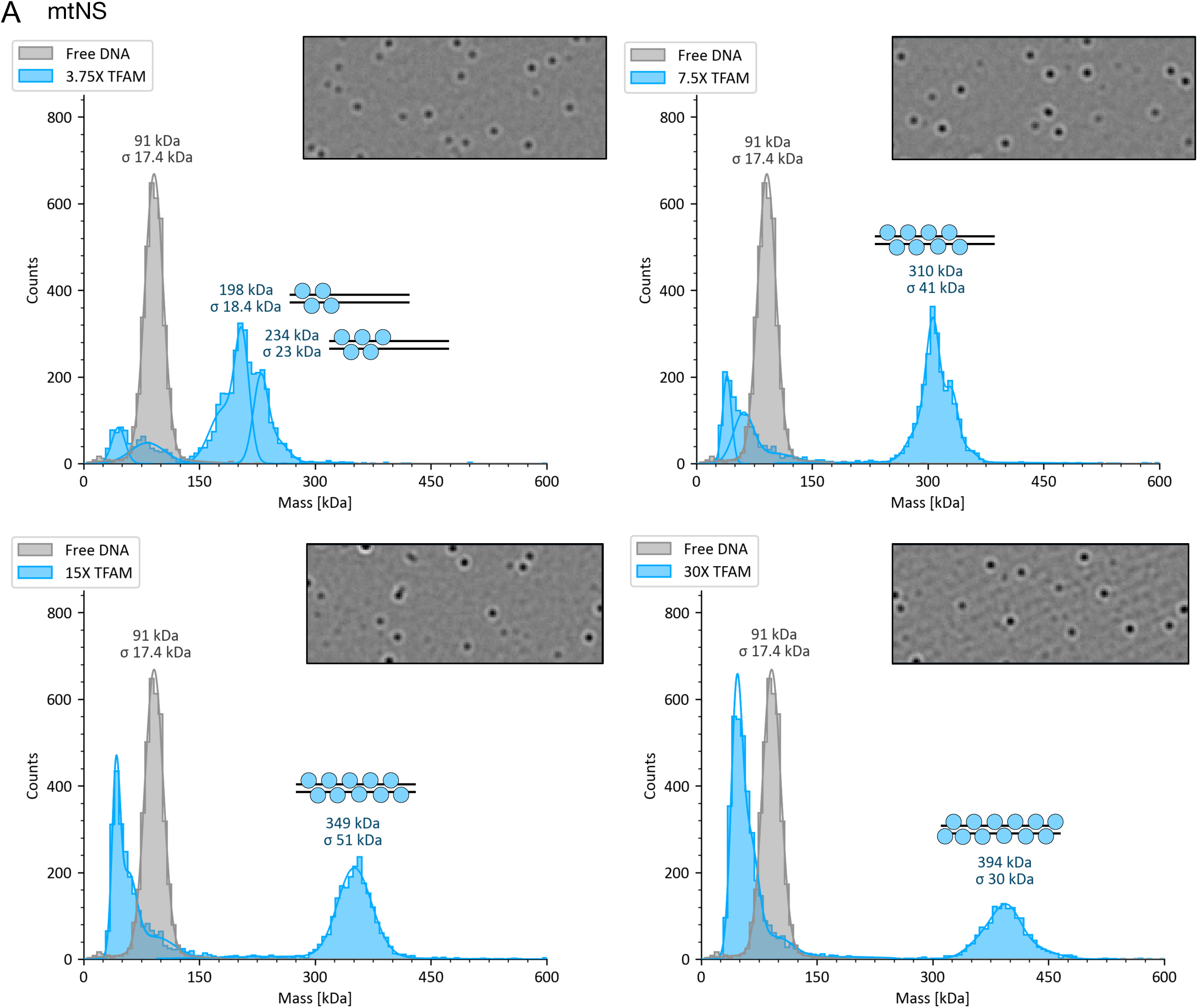
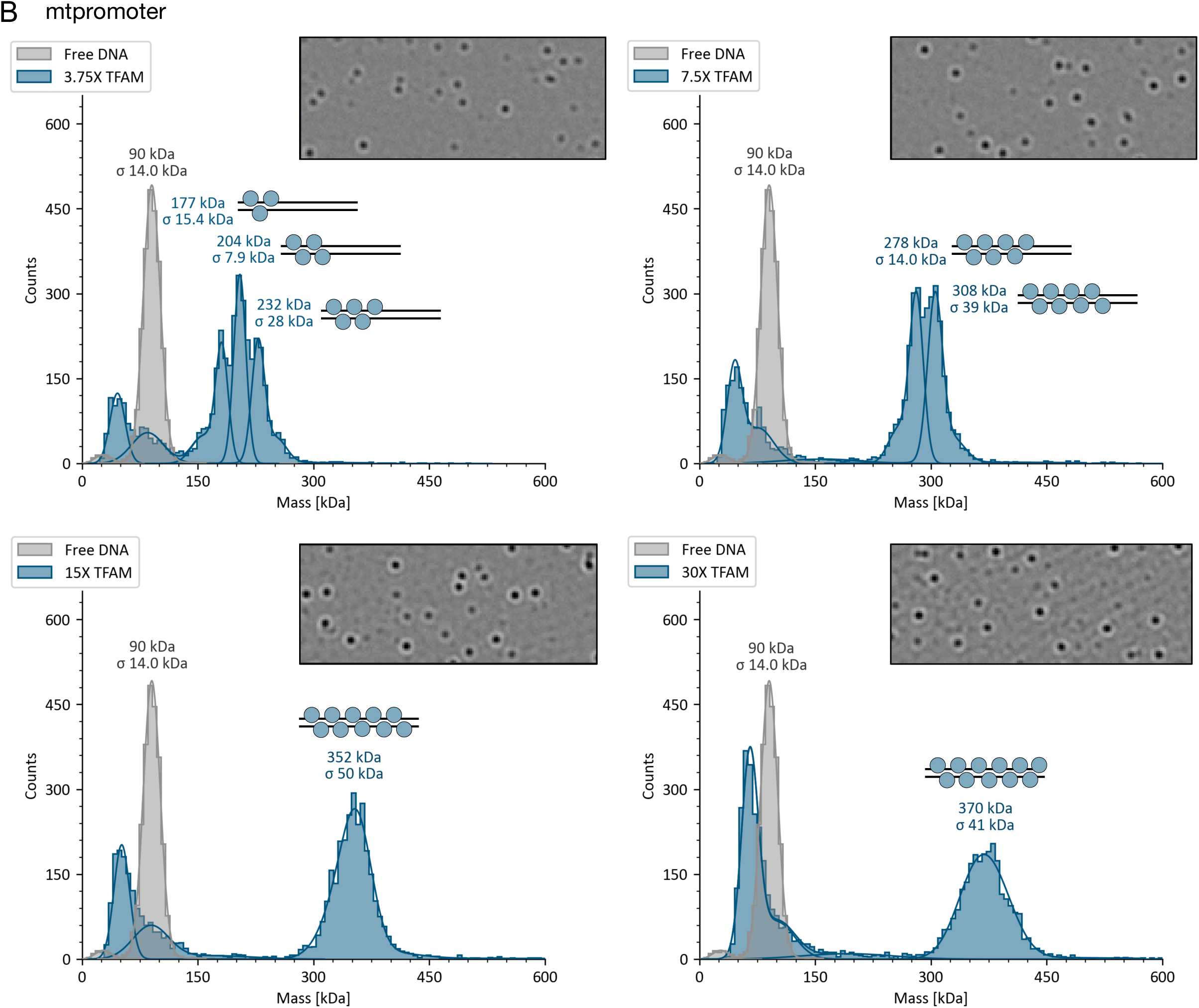
Mass photometry data for full titration with mitochondrial DNA-derived DNAs. **A.** TFAM titration into 160 bp mtNS DNA. **B.** TFAM titration into 160 bp mtpromoter DNAs. Cartoon near each peak represents the number of TFAMs bound in each detected complex.

**Figure S5.**
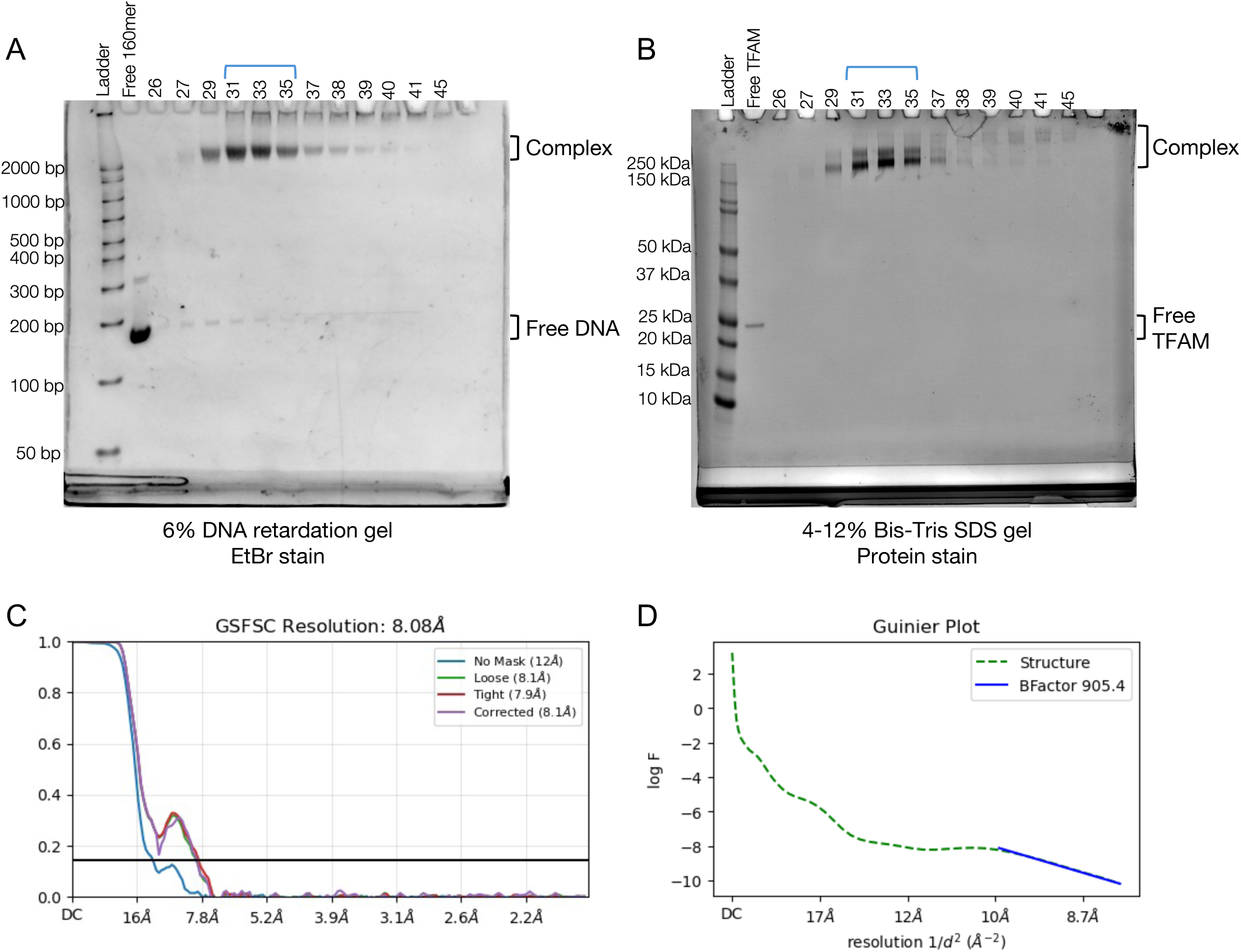
Sample preparation and refinement data for crosslinked TFAM-160mer complex. **A.** Fractions analyzed on 6% DNA retardation gels (Invitrogen) stained with Ethidium Bromide **B.** Fractions analyzed on SDSPAGE gels stained with Blazin’ blue. Pooled fractions are indicated by the blue bracket. **C.** Gold standard FSC curve for the density map from homogenous refinement. **D.** Guinier plot from homogenous refinement used to estimate B-factor of the density map.

**Figure S6.**
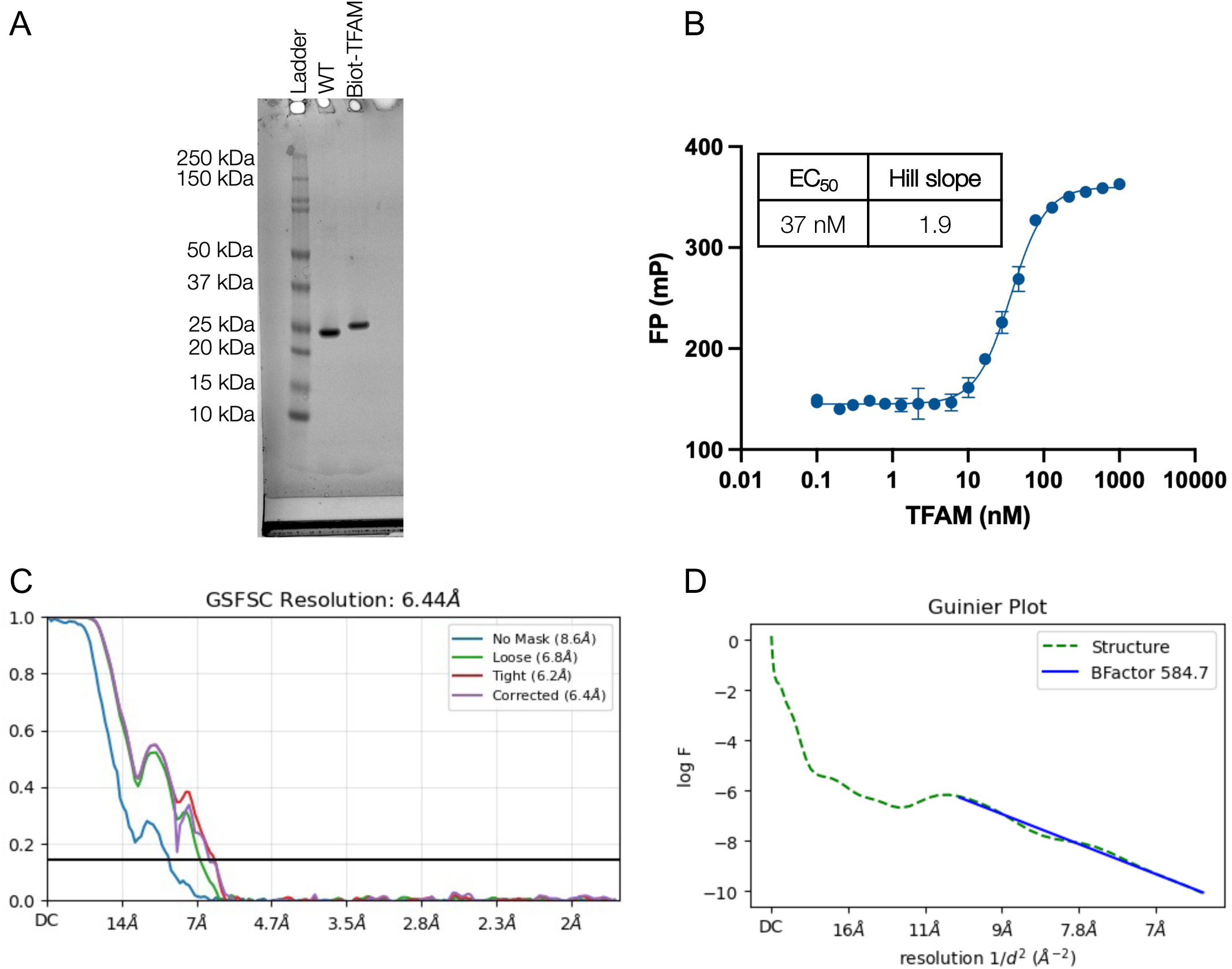
TFAM biotinylation for streptavidin monolayer affinity grids and refinement data for native TFAM-160mer complex. **A.** Verification of TFAM modification using 4-12% SDS-PAGE stained with Ethidium Bromide. **B.** Fluorescence Polarization with biotinylated TFAM and Alexa Fluor 488-labeled 160 bp DNA showing a similar binding affinity to wild type TFAM. **C.** Gold standard FSC curve for the density map from homogenous refinement. **D.** Guinier plot from homogenous refinement used to estimate B-factor of the density map.

**Figure S7.**
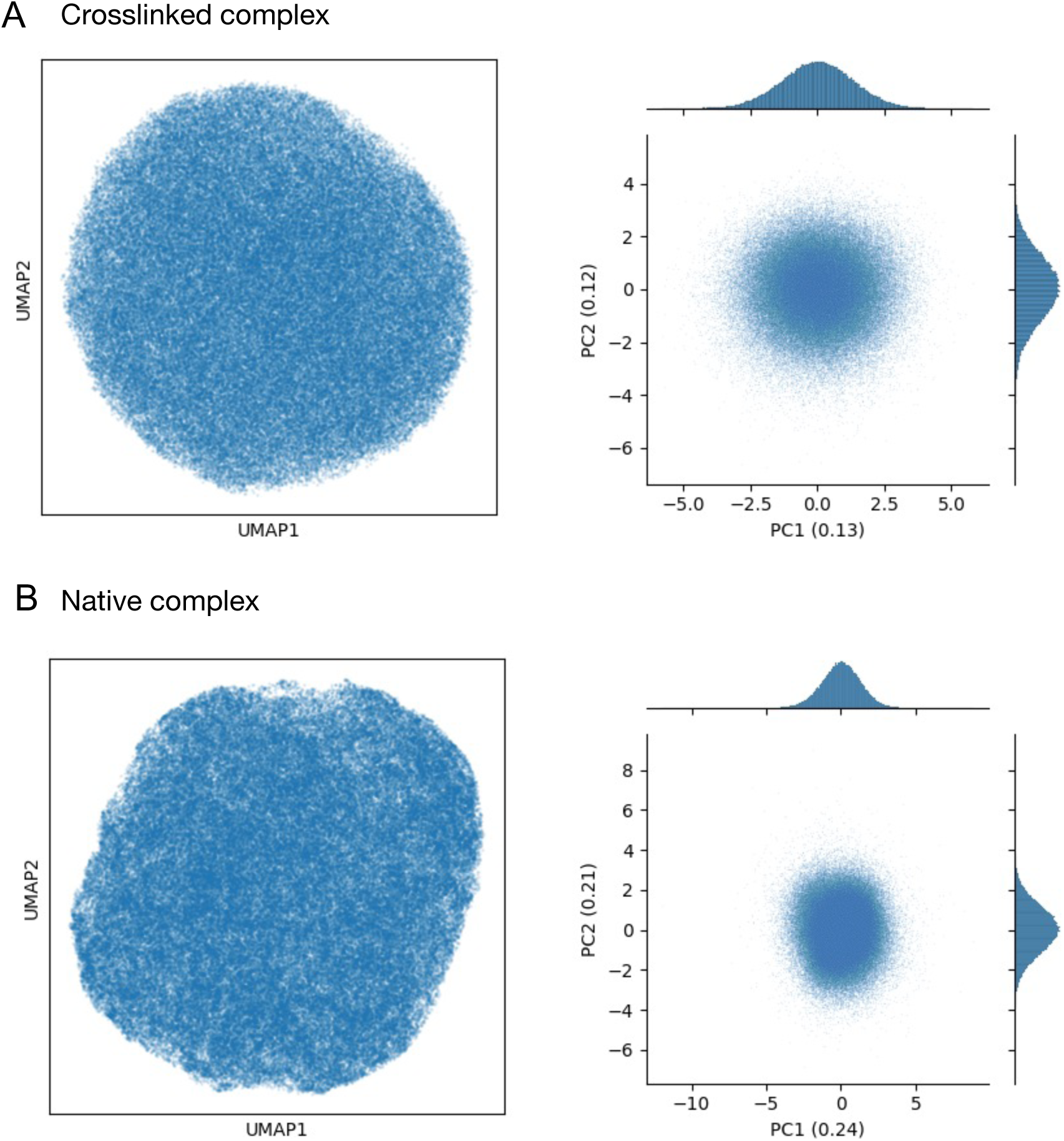
cryoDRGN analysis of native and crosslinked TFAM-160mer complexes. **A.** UMAP visualization (left) of the latent variable distribution and Principal Component Analysis (PCA) projection (right) of latent space encodings after training on a 10-dimensional latent variable model in cryoDRGN for the crosslinked complex. **B.** UMAP visualization (left) of the latent variable distribution and Principal Component Analysis (PCA) projection (right) of latent space encodings after training on a 10-dimensional latent variable model in cryoDRGN for the native complex.

